# PLD3 is a neuronal lysosomal phospholipase D associated with β-amyloid plaques and cognitive function in Alzheimer’s disease

**DOI:** 10.1101/746222

**Authors:** Alex G. Nackenoff, Timothy J. Hohman, Sarah M. Neuner, Carolyn S. Akers, Nicole C. Weitzel, Alena Shostak, Shawn Ferguson, David A. Bennett, Julie A. Schneider, Angela L. Jefferson, Catherine C. Kaczorowski, Matthew S. Schrag

**Author notes:** Corresponding author: Matthew S. Schrag, M.D., Ph.D., 6158 MRBIII, Department of Neurology, Vanderbilt University School of Medicine, Nashville TN, 37240.

## Abstract

Phospholipase D3 (PLD3) is a protein of unclear function that structurally resembles other members of the phospholipase D superfamily. A coding variant in this gene confers increased risk for the development of Alzheimer’s disease (AD), although the magnitude of this effect has been controversial. Because of the potential significance of this obscure protein, we undertook a study to determine whether PLD3 is relevant to memory and cognition in sporadic AD, to observe its distribution in normal human brain and AD-affected brain, to describe its subcellular localization, and to evaluate its molecular function. PLD3 mRNA levels in the human pre-frontal cortex inversely correlated with β-amyloid pathology severity and rate of cognitive decline in 531 participants enrolled in the Religious Orders Study and Rush Memory and Aging Project. PLD3 levels across genetically diverse BXD mouse strains and strains crossed with 5xFAD mice correlated strongly with learning and memory performance in a fear conditioning task. In human neuropathological samples, PLD3 was primarily found within neurons and colocalized with lysosome markers (LAMP2, progranulin, and cathepsins D and B). This colocalization was also present in AD brain with prominent enrichment on lysosomal accumulations within dystrophic neurites surrounding β-amyloid plaques. This pattern of protein distribution was conserved in mouse brain in wild type and the 5xFAD mouse model of cerebral β-amyloidosis. We discovered PLD3 has phospholipase D activity in lysosomes. A coding variant in PLD3 reported to confer AD risk significantly reduced enzymatic activity compared to wild-type PLD3. In summary, this study identified a new functional mammalian phospholipase D isoform which is lysosomal and closely associated with both β-amyloid pathology and cognition.

## Introduction

Alzheimer’s disease (AD) is the most common form of dementia and an increasing societal and economic burden. Multiple gene variants were recently discovered in the Phospholipase D3 (PLD3) gene which conferred increased risk for late onset AD^1^. Phospholipase D enzymes (PLDs) form a superfamily that are present in viruses, bacteria, and plants all the way up to mammalian cells. PLDs play an important role in the conversion primarily of phosphatidylcholine to phosphatidic acid and choline, which has implications for membrane dynamics and cell signaling, among other important cellular processes^2^. The phosphodiesterase domains of PLDs contain two highly conserved catalytic regions, HxKxxxxD motifs (hereafter HKD motif). There are a few non-canonical functions of PLDs which include nuclease activity and cardiolipase activity, both of which are mediated by substrate binding at the HKD motif^3,4^. Phospholipase D1 and D2 (PLD1 and PLD2), are confirmed to be enzymatically active mammalian PLD isoforms and are druggable proteins^2,5^. PLD3 was identified as a PLD based upon homology as it contains two HKD motifs, but has been informally reported to lack PLD activity^6^. The PLD3 gene variants associated with increased late onset AD risk were initially linked to β-amyloid precursor protein (APP) processing^1^ but this could only be replicated in overexpression conditions^7^. Moreover, a number of subsequent reports failed to replicate the association of the highest LOAD risk PLD3-V232M coding variant, although this may have been due in part to the rarity of the coding variant^8–11^. In a later meta-analysis, the PLD3-V232M contributed to AD risk but had a smaller effect than initially reported, comparable in magnitude to the apolipoprotein E-ε4 allele^12^. The V232M variant is associated with decreased PLD3 expression and is adjacent to an HKD motif, suggesting PLD3 hypofunction may drive or modulate AD pathology. Because the molecular function of this protein in the brain is unclear, it is not known whether this variant affects protein function. Additionally, because the variant is rare (∼1% of affected patients)^1^, the relevance of PLD3 to AD processes at a population level is yet unknown.

In all, there has been little resolution to the controversy whether PLD3 is a legitimate AD-risk gene, in part due to the limitations of studying rare coding gene variants. In this study, we aim to address the impact of PLD3 upon AD-related biological and cognitive pathologies leveraging generalizable population level datasets as well as determining the molecular function of PLD3.

### Variance in human PLD3 mRNA levels correlated with β-amyloid deposition and cognition in aged and AD participants

We evaluated whether stratifications in human PLD3 gene expression in the brain correlate with AD risk. The Religious Orders Study (ROS) and Rush Memory and Aging Project (MAP) enrolled older adults without dementia who agreed to annual clinical evaluations and brain donation^13–15^. A total of 531 ROS/MAP participants had mRNA expression levels measured from frozen sections of dorsolateral prefrontal cortex, as previously described^16^, along with autopsy measures of neuropathology and premorbid longitudinal neuropsychological test performance. Participants were well educated, predominantly female, predominantly non-Hispanic white, and on average 89 (±7) years of age at death (**Supplemental Table 1**). Detailed description of neuropsychological measures, RNA extraction and processing, neuropathological evaluation, and statistical analyses are presented in **Online Methods**.

In linear regression models covarying for age at death, sex, and postmortem interval, higher prefrontal cortex expression of *PLD3* was associated with lower β-amyloid plaque burden measured via immunolabeling, explaining 3% of the variance beyond covariates (**Figure 1**; PLD3: β=-0.004, p=1.7×10^-6^). β-amyloid plaque burden remained significantly associated when covarying for clinical diagnosis (p=1.6×10^-5^). Similarly, higher prefrontal cortex expression of *PLD3* was associated with neurofibrillary tangle burden quantified by silver stain (β=-0.0006: p=0.03) and IHC (β=-0.002, p = 0.03), however in both tangle models, inclusion of *in vivo* clinical diagnosis attenuated *PLD3* associations (p>0.07). When covarying for age at death, sex, and post mortem interval, lower levels of *PLD3* were observed among AD cases compared to cognitively normal participants (β=-11.9, p=0.03). PLD3 levels were not associated with cerebral atherosclerosis (**Supplementary Figure 1).**

**Figure 1.**
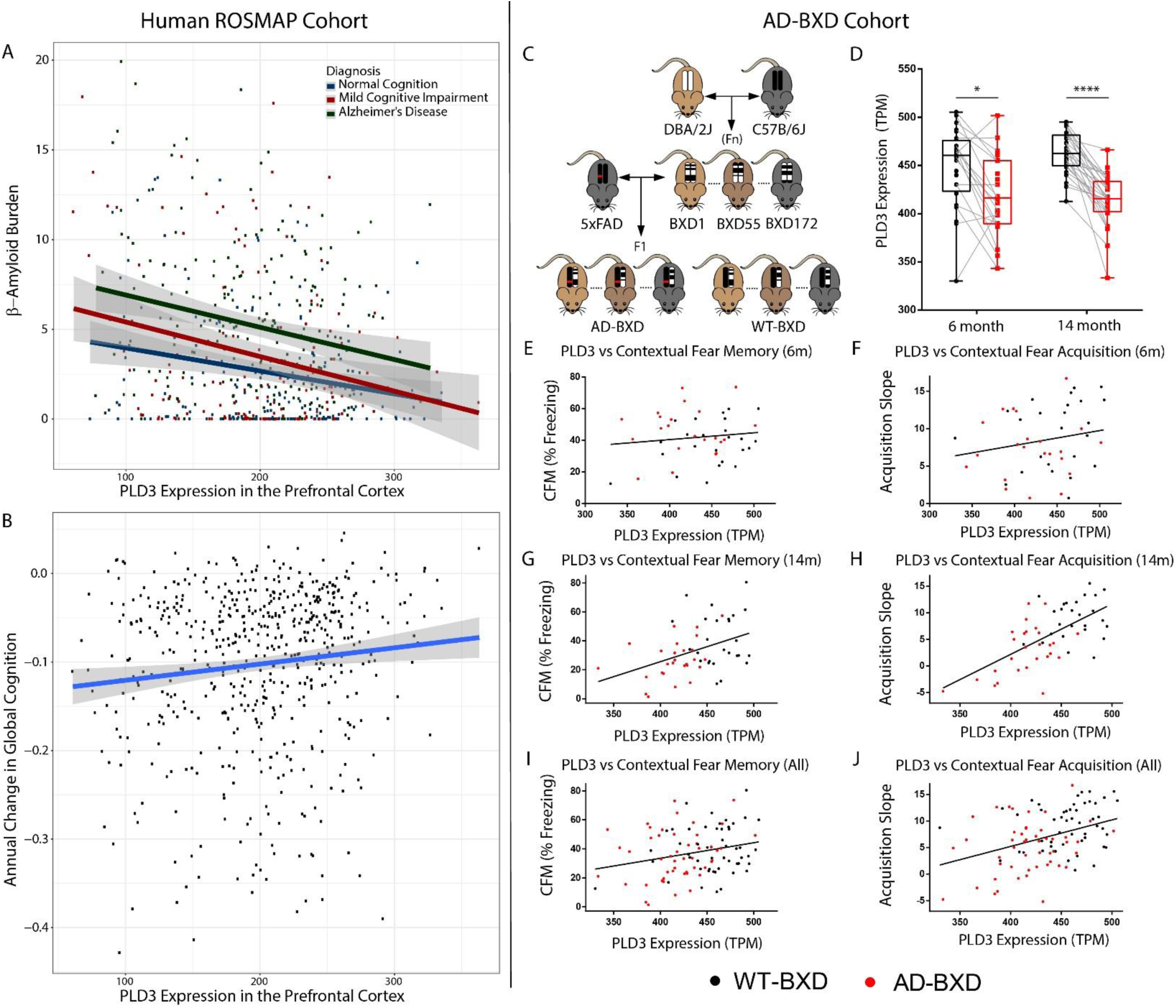
Variation in neuronal PLD3 expression predicts cognitive decline. **A-B.** Human ROS/MAP hippocampal RNA-Seq derived normalized PLD3 mRNA counts (in transcripts per million, TPM) **A.** Regression analysis reveals across all clinical diagnoses, higher levels of PLD3 appear protective with respect to lower β-amyloid plaque counts in prefrontal cortex. **B.** Across all clinical diagnoses, higher levels of PLD3 correlate with slower rates of cognitive decline. **C-J.** AD-BXD mimic human genetic diversity with respect to variation of PLD3 expression and subsequent learning and memory deficits. Mice underwent contextual fear testing (n=636), of which a subset was euthanized and brain tissue subjected to RNA-Seq (n=133). Dots represent averaged behavioral performance within BXD background strain and PLD3 transcript count (n=100 observations). **C.** AD mouse model 5xFAD (hemizygous) crossed with genetically diverse BXD model^17^. Littermate controls are generated via this F1 cross due to the hemizygosity of the 5xFAD transgene. **D.** PLD3 hippocampal mRNA expression (displayed in normalized transcripts per million (TPM)) is widely stratified across BXD background and is reduced in AD-BXD mice at both 6 and 14 months (Two-Way ANOVA, *p<0.05 & ****p<0.0001 following Bonferroni post-hoc tests). Gray connecting lines indicate shared sex and BXD background strain for visual comparison. **E-J.** Contextual fear conditioning behavioral performance regarding fear memory recall (**E, G, I**) and fear learning acquisition speed (**F, H, J**) plotted against the group averaged PLD3 mRNA transcript count. Pooled cohort of mice display positive correlation with hippocampal PLD3 mRNA expression and fear memory recall (**I**) and fear learning acquisition speed (**J**), with the correlational slope more strongly affected by age in memory recall (**G vs E**) and acquisition speed (**H vs F**).

In cognitive analyses, high prefrontal cortex expression of *PLD3* was associated with a slower rate of global cognitive decline when adjusting for age at death, sex, postmortem interval, and the interval between the final neuropsychological assessment and death (β=0.0002, p=0.02). Additional cognitive results are presented in **Supplementary Figure 2**.

It is also notable that we observed a subtle sex difference in *PLD3* associations. Prefrontal cortex levels of *PLD3* interacted with sex on β-amyloid plaque density measured with IHC (β=0.004, p=0.02) and on cross-sectional global cognitive performance at the final visit before death (β=-0.004, p=0.03). In both cases, *PLD3* associations were stronger in female than male participants (**Supplemental Figure 3**).

In replication samples from Mayo Clinic and Mt. Sinai accessed through the Accelerating Medicines Partnership AD project (AMP-AD) (**Supplemental Table 1)**, we observed lower levels of *PLD3* expression among AD cases compared to controls in the parahippocampal gyrus (p=0.0001), the cerebellum (p=0.003), and the temporal cortex (p=0.03) (**Supplemental Figure 4**).

### PLD3 mRNA levels are lower in AD-BXD mice and correlated with learning and memory performance

The BXD mouse model was created by crossing a wild-type mouse with C57BL/6J background with a DBA/2J mouse to produce a family of inbred sub-strains that capture a range of genetic diversity. AD-BXD mice were produced by crossing the resulting BXD strains with 5xFAD hemizygotes, an animal model containing five pro-β-amyloidosis mutations^17^ (**Figure 1C**). PLD3 mRNA levels were significantly lower in AD-BXD mice than their wild type BXD counterparts **(Figure 1D)**. Mice were subjected to a modified contextual fear memory test (as previously described^17^) due to its well-validated sensitivity to discern learning and memory impairments in multiple AD mouse models^18,19^. The first experimental day of measuring freezing behavior following repeated mild foot-shocks allows for the detection of learning acquisition rates, which differed among the WT-BXD and AD-BXD (**Figure 1F/H/J**). Additionally, the second experimental day measuring freezing behavior in the same environmental context without the presentation of foot-shocks allows for analysis of memory recall, which varied within WT-BXD and AD-BXD mice (**Figure 1E/G/I**). Following behavioral experimentation at the respective time points, a WT and 5xFAD mouse from each strain was euthanized to analyze mRNA levels in hippocampus via RNA-Seq. We modeled strain-averaged behavioral performance against PLD3 mRNA transcript counts and found that PLD3 levels (adjusting for age and sex) predicted learning acquisition slope (β=0.06, p=2.8×10^-6^, explaining 6% of the variance) (**Figure 1F/H/J**). When adjusting for age and sex, PLD3 levels were a significant predictor of fear memory recall (β=0.09, p=0.03) (**Figure 1E/G/I)**. When including genotype (β-amyloid state) and BXD background strain, PLD3 remained predictive of learning acquisition speed (β=0.04, p=0.003) but not when modeling fear memory recall (β=0.085, p=0.08). These findings are unsurprising as the main effect for the AD-BXD behavioral stratification was observed upon learning acquisition slope rather than memory recall, as memory recall is more strongly sensitive to β-amyloid burden^17,20^. Collectively, these human and animal data suggest that PLD3 expression has a direct impact upon cognition in the aged brain.

### PLD3 is a lysosomal protein and enriched around β-amyloid plaques

While most early reports suggested PLD3 was an endoplasmic reticulum-localized protein^21^, a few studies suggested it may, at least partially, localize to the lysosome. Most of these claims were based on studies of overexpression of the protein^22,23^. Because of contradictory reports regarding the subcellular localization of PLD3, we sought to better resolve the subcellular localization of PLD3 with attention to endogenous context-relevant tissue. In human brain tissue, PLD3 staining in neuronal cell bodies colocalized highly with Cathepsin B (mean Pearson correlation coefficient 0.84±0.07 for neurological controls, 0.86±0.04 for AD) and other lysosomal (LAMP1 and Cathepsin D in **Supplemental Figure 6**). In AD, β-amyloid plaques are surrounded by enlarged axon segments, termed dystrophic neurites, which are filled with lysosome-like organelles^24^ (**Figure 2**). In dystrophic neurites around β-amyloid plaques, PLD3 was enriched and strongly localized with lysosomal membrane markers (**Figure 2**), although the association with luminal markers Cathepsin B/D was attenuated (**Figure 2**), consistent with a prior report that lysosomes in dystrophic neurites are deficient in luminal proteases^24^. These data confirm that even around β-amyloid plaques, PLD3 remains on lysosomes. Similarly, in brain tissue from wild-type and 5xFAD mice PLD3 was strongly colocalized with markers of lysosomes (Pearson coefficient: 0.72±0.22 for WT, 0.65±0.10 for 5xFAD). The AD mouse model 5xFAD recapitulates this phenotype (**Figure 2**). Furthermore, in 5xFAD PLD3 did not colocalize with β-amyloid (Pearson correlation coefficient 0.13±0.10), but was enriched on lysosome-like organelles within dystrophic neurites surrounding β-amyloid plaques. Finally, to confirm that PLD3 is a lysosomal protein, lysosomes were isolated from non-transfected HeLa cells by magnetic fractionation after “pulse chase” loading of dextran-coated iron-oxide nanoparticles (**Supplemental Figure 7**). PLD3 was highly enriched in the lysosomal fraction and accounted for the vast majority of the PLD3 in the cells (**Figure 3B**), confirming that PLD3 is primarily a lysosomal protein.

**Figure 2.**
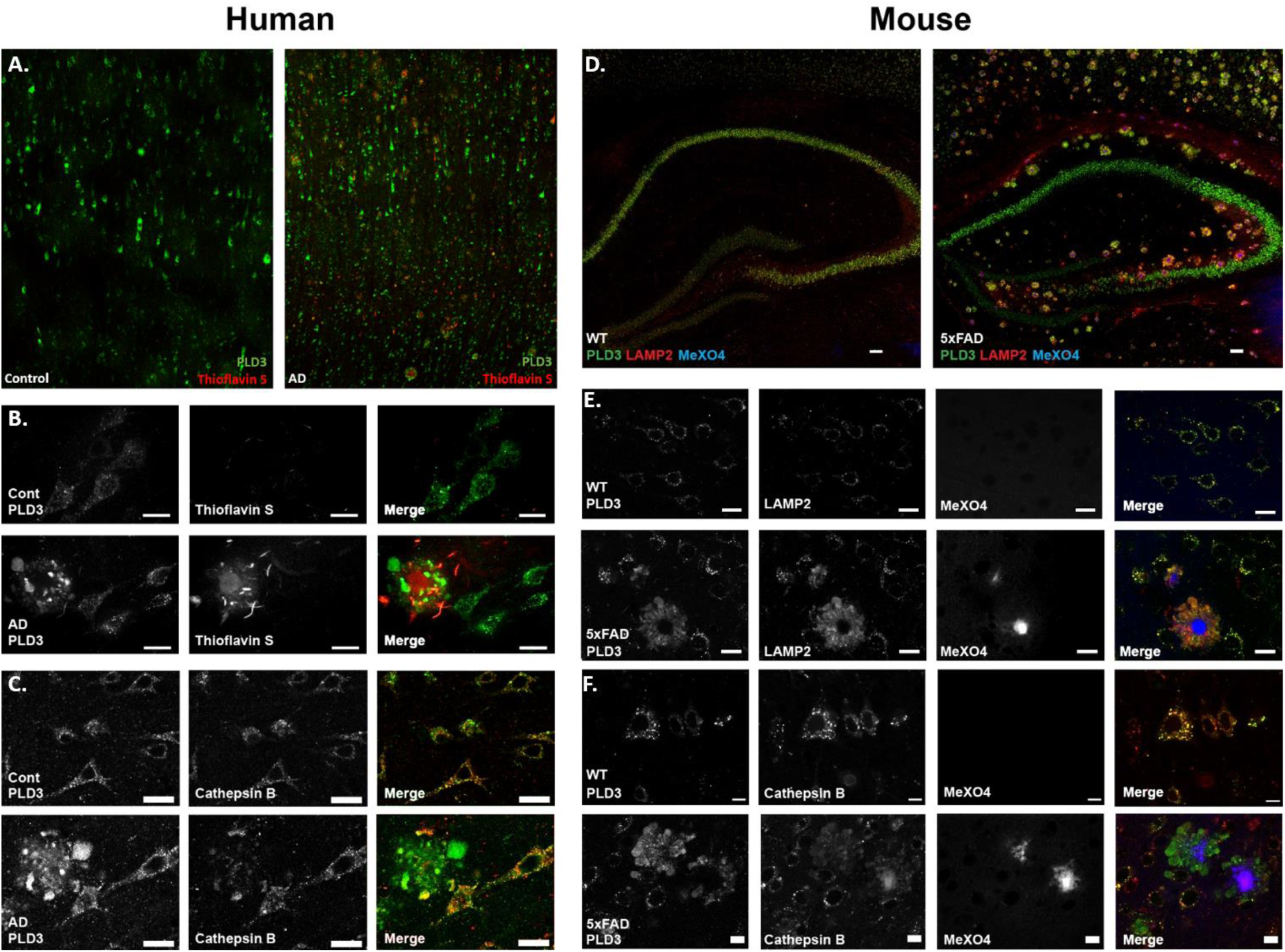
PLD3 is enriched in lysosomes surrounding parenchymal β-amyloid plaques in AD tissue. **A.** Low magnification images of temporal lobe cortex from neurological control and AD brain demonstrated a neuronal pattern of staining against PLD3 (green) and consistent accumulation around β-amyloid plaques. **B.** 40x magnification images of the same immunohistochemistry shows PLD3 is in a punctate staining pattern within neuronal cell bodies and enriched around β-amyloid plaques. **C.** Co-staining with lysosomal marker cathepsin B confirmed PLD3 was primarily lysosomal and enriched in dystrophic neurites in AD brain. **D.** In 5xFAD mice, PLD3 was similarly enriched around every β-amyloid plaque and strongly colocalized with LAMP2. **E.** 40x magnification of WT and 5xFAD tissue demonstrated strong co-localization of PLD3 with lysosomal membrane marker LAMP2 (Pearson coefficient: 0.84±0.07 for WT, 0.86±0.04 for 5xFAD) and strong staining in dystrophic neurites around β-amyloid plaques, which are stained blue with methoxy-X04. **F.** The lysosomal lumenal protease cathepsin B is similarly co-localized with PLD3 in both normal and diseased neurons.

**Figure 3.**
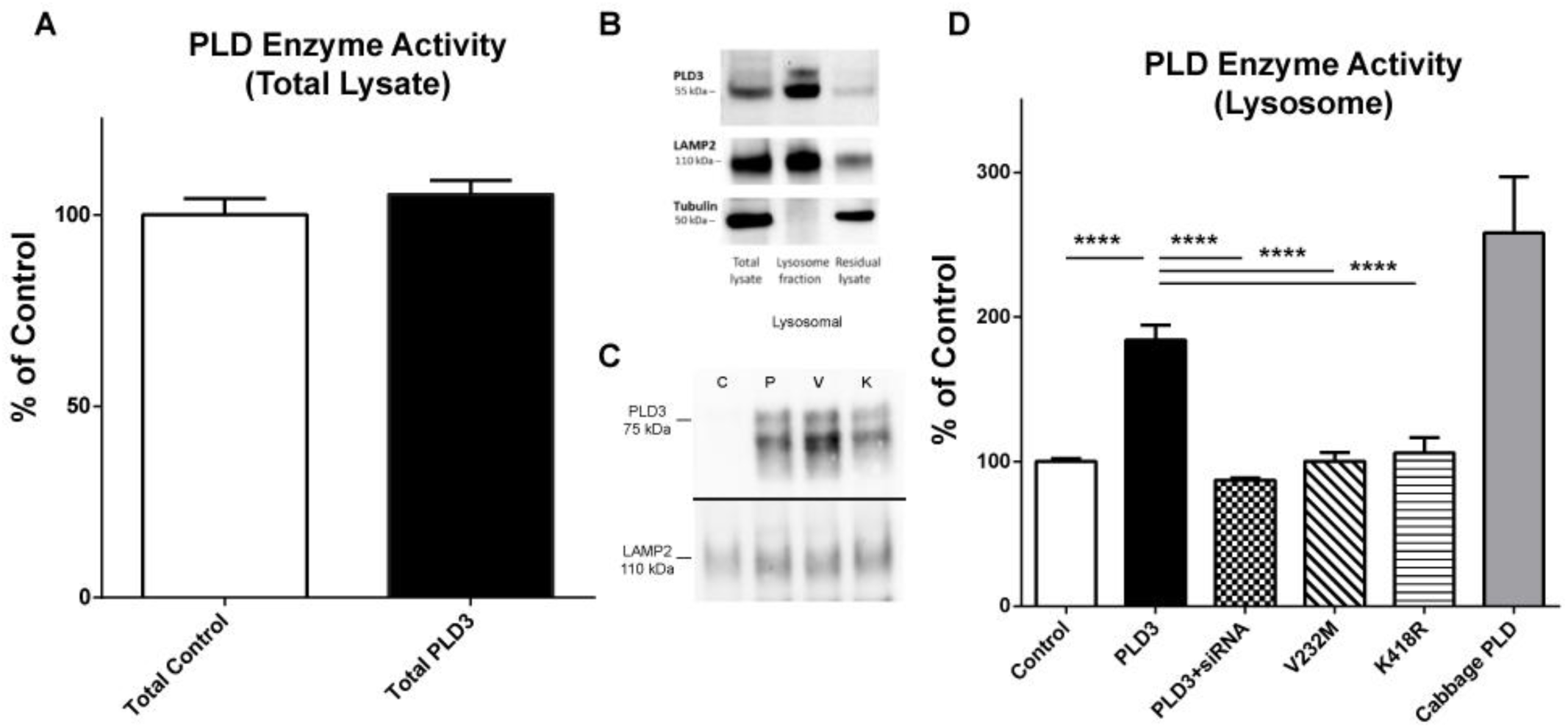
PLD3 displays phospholipase D activity in intact isolated lysosomes. **A.** Overexpression of PLD3 does not produce increased phospholipase D activity in total lysate of NSC cells compared to untransfected NSC total lysate. N=4 biological/assay replicates. **B.** Lysosomes isolated following transfection of PLD3 via iron dextran nanoparticles and magnetic affinity columns display enrichment for lysosomal marker LAMP2 and expected lack of cytoskeletal marker tubulin. Lysosomal fraction is enriched with PLD3, further indicating that PLD3 is a lysosomal protein. **C.** Induction of AD associated PLD3 variant V232M and HKD mutation K418R does not impact total PLD3 expression (data not shown) or routing to the lysosome (representative blot shown), indicated by Western Blot. Samples isolated via lysosomal isolation detailed in the Methods section. **D**. Data derived from lysosomes isolated from transfected NSC cells, each condition containing n=4 biological/assay replicates per condition and n>16 sample replicates per condition. Phospholipase D (PLD) activity measured by the Amplex Red PLD assay with negative (control lysosomes) and positive (Cabbage PLD) controls in each assay run. Data analyzed via One-Way ANOVA. ****p<0.0001 following Bonferroni post-hoc tests. Lysosomes expressing PLD3 display significant increase in PLD activity compared to control lysosomes, which is near the magnitude of PLD activity seen with purified cabbage PLD but significantly different (p<0.001). Upon cotransfection of siRNA against PLD3 alongside PLD3 transfection, these lysosomes display no apparent PLD activity. PLD3-K418R transfected lysosomes, which has a mutated putative lysine in its HKD domain, displays significantly reduced PLD activity compared to PLD3 transfected lysosomes. Furthermore, human AD-risk associated variant PLD3-V232M displays no significant PLD activity, and displays a significant reduction in PLD activity compared to PLD3.

### PLD3 has phospholipase D activity in intact lysosomes

There was no phospholipase D activity above baseline in total lysate of PLD3 transfected mammalian cells (**Figure 3A**). However, when we isolated lysosomes from PLD3-transfected NSC34 cells (a murine neuroblastoma line) to assess PLD activity, we found that PLD3 containing lysosomes had significantly increased phospholipase D activity compared to non-transfected lysosomes (**Figure 3D**). The specificity of the increase in PLD activity was evaluated by co-transfecting a validated siRNA against hPLD3 (validated in **Supplemental Figure 8**). Co-transfection with anti-PLD3 siRNA reduced the phospholipase D activity to that of untransfected lysosomes (**Figure 3D**). The specificity of PLD3 activity via siRNA knockdown was replicated in a human cell line (**Supplemental Figure 9**). To further confirm that the phospholipase D activity originated from the conserved HKD domain, we introduced a mutation in the second HKD domain, K418R, which completely ablated the enzymatic activity of PLD3 (**Figure 3D**). A plasmid containing a K203R mutation in the first HKD domain did not efficiently express full length modified protein (data not shown). Finally, we found the human AD-associated V232M variant lacked PLD activity (**Figure 3D**). These data suggest PLD3 is a functional phosopholipase D in lysosomes. Additionally, PLD3 transfection did not appear to upregulate PLD1 or PLD2 in the lysosome (**Supplemental Figure 10**), which are known to be functional PLD enzymes. In a cell-free assay, affinity purified His-tagged full-length PLD3 isolated from mammalian cell lysates did not have detectable PLD activity under a range of conditions and via various detection methodologies (data not shown). The presence of PLD3 activity in intact lysosomes and our inability to reproduce this activity in purified cell-free systems suggests a functional contribution of organelle-specific post-translational modifications, activating protein binding partners, or some other feature of the lysosomal microenvironment for phospholipase D activity of PLD3.

## Discussion

Rare PLD3 coding gene variants conferring an increased risk of late onset AD were discovered in 2014 but have remained controversial in large part due to the unknown function of PLD3 and unclear molecular impact of the rare variants^1^. Here, we leveraged multimodal datasets that reflect the role of PLD3 at a population level in both a large, longitudinal human study and in a genetically diverse murine model to establish a functional connection between PLD3 expression and AD-related cognitive impairment. In humans, we found a protective association between increased PLD3 expression and β-amyloid plaque burden. Increased PLD3 expression correlated with reduced rate of cognitive decline in the ROSMAP cohort and learning and memory in the AD-BXD mice (**Figure 1**). These results advance our understanding of the functional connection between PLD3 expression and cognition to establish that PLD3 may be relevant to AD at a population level.

Endolysosomal dysfunction in AD has become an important research focus, in part because dystrophic neurites are a consistent pathological feature of AD made up of enlarged axon segments around β-amyloid plaques clogged with dysfunctional lysosomes^24,25^. Here, we report PLD3 is highly concentrated in these abnormal neuronal lysosomes surrounding β-amyloid plaques (**Figure 2**), corroborating earlier reports of PLD3 upregulation around β-amyloid plaques^26^ and that PLD3 can be routed to the lysosome^22^. The close and consistent association of PLD3 with β-amyloid neuropathology further strengthens the relevance of PLD3 to AD.

Finally, we evaluated the molecular function of PLD3. Because PLD3 is lysosomal at endogenous expression levels, we hypothesize that its activity likely requires an acidic pH or some other feature(s) of the lysosomal microenvironment. Consistent with initial reports^6^, we did not detect a change in phospholipase D activity in total cell lysates after overexpressing PLD3, which is not surprising given lysosomes account for well under 5% of the total lysate and several other PLD isoforms are present in the lysate. However, when we isolated lysosomes from transfected mammalian cell lines, we discovered markedly enhanced PLD activity after transfecting PLD3, which was ablated when an anti-PLD3 siRNA was co-transfected or a mutation in the putative active site was introduced. Using this assay, we demonstrated that the AD-associated PLD3 variant, V232M, lacked PLD activity. This discovery strengthens the case that PLD3-V232M is a bonafide AD risk gene. Moreover, these results give us confidence that PLD3 is a mammalian PLD isoform.

An alternative molecular function of PLD3 has been proposed. A recent report argued that both PLD3 and PLD4 have 5’ exonuclease activity and implicated this activity in the function of antigen presenting cells in the context of autoimmune diseases^3^. PLD4 variants have been linked to autoimmune disease, particularly rheumatoid arthritis^27–29^, but there is no such association with PLD3. Moreover, we found PLD3 was nearly exclusively expressed in neurons in the brain and was essentially absent from microglia. PLD3 knock-out mice have been reported to have increased lysosomal size in neurons^7^, and an alteration in neuronal lysosomal membrane dynamics (fission or fusion) is more consistent with PLD activity than with exonuclease activity. Our inability to reproduce the PLD activity of PLD3 in a purified cell-free system is a limitation but not unexpected. Both PLD1 and PLD2 require various activating factors for *in vitro* activity^30^, it is therefore highly likely that PLD3 requires similar factors which are currently unknown but are presumably present in the lysosomal microenvironment. It may ultimately be necessary to decipher the structure of PLD3 in greater detail than currently available to confirm the primary molecular function. Importantly, however, whether PLD3 is a PLD or 5’ exonuclease, the HKD domains appear to be enzymatically active which raises the probability that an inactivating mutation in PLD3 is truly pathogenic.

The enrichment of PLD3 in dystrophic neuronal lysosomes surrounding β-amyloid plaques in AD is, at first glance, incongruent with our human and mouse population level data indicating higher levels of PLD3 are proadaptive with respect to β-amyloid plaque burden and cognition. While it is possible that PLD3 enrichment in dystrophic neurites represents a compensatory cellular response to plaque formation, we hypothesize that PLD3 is functionally inactive in these sites. Lysosomes in dystrophic neurites appear to be functioning sub-optimally based on our observation that they are lacking in lysosomal proteases^24^ indicated by a deficiency in luminal protease markers Cathepsin B/D (**Figure 2**). Given this functional defect, it is reasonable to hypothesize that they might also fail to acidify, and if PLD3 required acidic pH to be active, it could be functionally inactive in dystrophic neurites. This highlights the need for further detailed studies to define the physiology of lysosomes within dystrophic neurites. It may be possible to interrogate dystrophic neurites *ex vivo* via organotypic tissue culture^31^ using tissue from the 5xFAD mouse or other AD mouse models. Together, these data warrant further investigation towards the potential protective effect of enhancing PLD3 activity to disrupt AD pathogenesis.

## References

1. Cruchaga, C. et al. Rare coding variants in the phospholipase D3 gene confer risk for Alzheimer’s disease. Nature 505, 550–554 (2013).

2. Brown, H. A., Thomas, P. G. & Lindsley, C. W. Targeting phospholipase D in cancer, infection and neurodegenerative disorders. Nat Rev Drug Discov 16, 351–367 (2017).

3. Gavin, A. L. et al. PLD3 and PLD4 are single stranded acid exonucleases that regulate endosomal nucleic acid sensing. Nat Immunol 19, 942–953 (2018).

4. Ipsaro, J. J., Haase, A. D., Knott, S. R., Joshua-Tor, L. & Hannon, G. J. The structural biochemistry of Zucchini implicates it as a nuclease in piRNA biogenesis. Nature 491, 279–283 (2012).

5. Saito, M. & Kanfer, J. Solubilization and properties of a membrane-bound enzyme from rat brain catalyzing a base-exchange reaction. Biochem. Biophys. Res. Commun. 53, 391–398 (1973).

6. Gonzalez, A. C. et al. PLD3 and spinocerebellar ataxia. Brain 141, e78–e78 (2018).

7. Fazzari, P. et al. PLD3 gene and processing of APP. Nature 541, E1–E2 (2017).

8. Hooli, B. V. et al. PLD3 gene variants and Alzheimer’s disease. Nature 520, E7–8 (2015).

9. Lambert, J.-C. et al. PLD3 and sporadic Alzheimer’s disease risk. Nature 520, E1 (2015).

10. Heilmann, S. et al. PLD3 in non-familial Alzheimer’s disease. Nature 520, E3–5 (2015).

11. van der Lee, S. J. et al. PLD3 variants in population studies. Nature 520, E2–3 (2015).

12. Engelman, C. D. et al. The effect of rare variants in TREM2 and PLD3 on longitudinal cognitive function in the Wisconsin Registry for Alzheimer’s Prevention. Neurobiology of Aging (2017). doi:10.1016/j.neurobiolaging.2017.12.025

13. Bennett, D. A., Schneider, J. A., Arvanitakis, Z. & Wilson, R. S. Overview and findings from the religious orders study. Curr Alzheimer Res 9, 628–645 (2012).

14. Bennett, D. A. et al. Overview and findings from the rush Memory and Aging Project. Curr Alzheimer Res 9, 646–663 (2012).

15. Bennett, D. A. et al. Religious Orders Study and Rush Memory and Aging Project. J. Alzheimers Dis. 64, S161–S189 (2018).

16. Mostafavi, S. et al. A molecular network of the aging human brain provides insights into the pathology and cognitive decline of Alzheimer’s disease. Nat. Neurosci. 21, 811–819 (2018).

17. Neuner, S. M., Heuer, S. E., Huentelman, M. J., O’Connell, K. M. S. & Kaczorowski, C. C. Harnessing Genetic Complexity to Enhance Translatability of Alzheimer’s Disease Mouse Models: A Path toward Precision Medicine. Neuron 101, 399–411.e5 (2019).

18. Puzzo, D., Lee, L., Palmeri, A., Calabrese, G. & Arancio, O. Behavioral assays with mouse models of Alzheimer’s disease: practical considerations and guidelines. Biochem. Pharmacol. 88, 450–467 (2014).

19. Webster, S. J., Bachstetter, A. D., Nelson, P. T., Schmitt, F. A. & Van Eldik, L. J. Using mice to model Alzheimer’s dementia: an overview of the clinical disease and the preclinical behavioral changes in 10 mouse models. Front Genet 5, (2014).

20. Neuner, S. M. et al. Systems genetics identifies modifiers of Alzheimer’s disease risk and resilience. bioRxiv 225714 (2017). doi:10.1101/225714

21. Osisami, M., Ali, W. & Frohman, M. A. A Role for Phospholipase D3 in Myotube Formation. PLOS ONE 7, e33341 (2012).

22. Gonzalez, A. C. et al. Unconventional Trafficking of Mammalian Phospholipase D3 to Lysosomes. Cell Reports 22, 1040–1053 (2018).

23. Demirev, A. V. et al. V232M substitution restricts a distinct O-glycosylation of PLD3 and its neuroprotective function. Neurobiology of Disease (2019). doi:10.1016/j.nbd.2019.05.015

24. Gowrishankar, S. et al. Massive accumulation of luminal protease-deficient axonal lysosomes at Alzheimer’s disease amyloid plaques. Proc Natl Acad Sci U S A 112, E3699–E3708 (2015).

25. Cataldo, A. M., Hamilton, D. J. & Nixon, R. A. Lysosomal abnormalities in degenerating neurons link neuronal compromise to senile plaque development in Alzheimer disease. Brain Res. 640, 68–80 (1994).

26. Satoh, J. et al. PLD3 is accumulated on neuritic plaques in Alzheimer’s disease brains. Alzheimer’s Research & Therapy 6, 70 (2014).

27. Okada, Y. et al. Meta-analysis identifies nine new loci associated with rheumatoid arthritis in the Japanese population. Nat. Genet. 44, 511–516 (2012).

28. Chen, W.-C. et al. rs2841277 (PLD4) is associated with susceptibility and rs4672495 is associated with disease activity in rheumatoid arthritis. Oncotarget 8, 64180–64190 (2017).

29. Akizuki, S. et al. PLD4 is a genetic determinant to systemic lupus erythematosus and involved in murine autoimmune phenotypes. Ann. Rheum. Dis. 78, 509–518 (2019).

30. Henage, L. G., Exton, J. H. & Brown, H. A. Kinetic analysis of a mammalian phospholipase D: allosteric modulation by monomeric GTPases, protein kinase C, and polyphosphoinositides. J. Biol. Chem. 281, 3408–3417 (2006).

31. Schommer, J., Schrag, M., Nackenoff, A., Marwarha, G. & Ghribi, O. Method for organotypic tissue culture in the aged animal. MethodsX 4, 166–171 (2017).

32. Oakley, H. et al. Intraneuronal beta-amyloid aggregates, neurodegeneration, and neuron loss in transgenic mice with five familial Alzheimer’s disease mutations: potential factors in amyloid plaque formation. J. Neurosci. 26, 10129–10140 (2006).

33. Schneider, J. A., Arvanitakis, Z., Bang, W. & Bennett, D. A. Mixed brain pathologies account for most dementia cases in community-dwelling older persons. Neurology 69, 2197–2204 (2007).

34. Mahoney, E. R. et al. Brain expression of the vascular endothelial growth factor gene family in cognitive aging and alzheimer’s disease. Mol Psychiatry 1–9 (2019). doi:10.1038/s41380-019-0458-5

35. Arvanitakis, Z. et al. The Relationship of Cerebral Vessel Pathology to Brain Microinfarcts. Brain Pathology 27, 77–85 (2017).

36. Arvanitakis Zoe, Leurgans Sue E., Barnes Lisa L., Bennett David A. & Schneider Julie A. Microinfarct Pathology, Dementia, and Cognitive Systems. Stroke 42, 722–727 (2011).

37. Boyle, P. A. et al. Cerebral amyloid angiopathy and cognitive outcomes in community-based older persons. Neurology 85, 1930–1936 (2015).

38. Buchman Aron S., Leurgans Sue E., Nag Sukriti, Bennett David A. & Schneider Julie A. Cerebrovascular Disease Pathology and Parkinsonian Signs in Old Age. Stroke 42, 3183–3189 (2011).

39. Love, S. et al. Development, appraisal, validation and implementation of a consensus protocol for the assessment of cerebral amyloid angiopathy in post-mortem brain tissue. Am J Neurodegener Dis 3, 19–32 (2014).

40. Schneider, J. A. et al. Relation of cerebral infarctions to dementia and cognitive function in older persons. Neurology 60, 1082–1088 (2003).

41. Schneider, J. A., Boyle, P. A., Arvanitakis, Z., Bienias, J. L. & Bennett, D. A. Subcortical infarcts, Alzheimer’s disease pathology, and memory function in older persons. Ann. Neurol. 62, 59–66 (2007).

42. Amador-Ortiz, C. et al. TDP-43 immunoreactivity in hippocampal sclerosis and Alzheimer’s disease. Ann. Neurol. 61, 435–445 (2007).

43. Wilson, R. S. et al. Temporal course and pathologic basis of unawareness of memory loss in dementia. Neurology 85, 984–991 (2015).

44. Wang, M. et al. The Mount Sinai cohort of large-scale genomic, transcriptomic and proteomic data in Alzheimer’s disease. Sci Data 5, 180185 (2018).

45. Allen, M. et al. Human whole genome genotype and transcriptome data for Alzheimer’s and other neurodegenerative diseases. Sci Data 3, 160089 (2016).

46. Allen, M. et al. Conserved brain myelination networks are altered in Alzheimer’s and other neurodegenerative diseases. Alzheimers Dement 14, 352–366 (2018).

47. Diettrich, O., Mills, K., Johnson, A. W., Hasilik, A. & Winchester, B. G. Application of magnetic chromatography to the isolation of lysosomes from fibroblasts of patients with lysosomal storage disorders. FEBS Lett. 441, 369–372 (1998).

