## Supplemental Materials & Methods for "PLD3 is a neuronal lysosomal phospholipase D associated with β-amyloid plaques and cognitive function in Alzheimer’s disease"

Supplemental Table 1: PLD3 expression levels by brain region in the Accelerating Medicines Partnership AD project (AMP-AD)

Supplemental Table 2: Human AD Brain Tissue Utilized for Neuropathological Immunohistochemistry

Supplemental Figure 1: PLD3 mRNA levels do not correlate with cerebral atherosclerotic Disease

Supplemental Figure 2: ROS/MAP Cognitive Sub-Measures

Supplemental Figure 3: Protective effect of PLD3 expression on β-Amyloid burden is stronger in women

Supplemental Figure 4: Multiple dataset analysis reveals PLD3 signal in AD risk models

Supplemental Figure 5: AD-BXD mouse performance on balance beam does not correlate with PLD3 expression

Supplemental Figure 6: PLD3 colocalizes with numerous markers of lysosomes in human brain

Supplemental Figure 7: Lysosomal isolation via iron oxide dextran coated nanoparticles

Supplemental Figure 8: Anti-PLD3 siRNA Deplete Total and Lysosomal PLD3 levels

Supplemental Figure 9: HeLa Derived Transfected Lysosomes Replicate PLD3 Specific Enzyme Function

Supplemental Figure 10: PLD3 transfection does not alter expression of PLD1 or PLD2

**Supplemental Table 1: PLD3 expression levels by brain region in the Accelerating Medicines Partnership AD project (AMP-AD)**

| ***PLD3* Differential Expression in Brain Tissues** | | | | | | | | | |
|---|---|---|---|---|---|---|---|---|---|
| **Model** | **Tissue** | **Study** | **logFC** | **CI.L** | **CI.R** | **AveExpr** | **t** | **P.Value** | **adj.P.Val** |
| Diagnosis | PHG | MSSM | -0.20499 | -0.30984 | -0.10015 | 8.664672 | -3.83868 | **0.000135** | **0.001454** |
| Diagnosis | CBE | MAYO | -0.17918 | -0.29537 | -0.06298 | 4.444553 | -3.0299 | **0.002577** | **0.010901** |
| Diagnosis | FP | MSSM | 0.119779 | 0.029162 | 0.210396 | 8.664672 | 2.595231 | **0.009652** | 0.151912 |

Adjusted P value corrected for all tests in each tissue (over 15,000). PHG=parahippocampal gyrus, CBE=cerebellum, FP=frontal pole.

**Supplemental Table 2. Neuropathology subjects' clinical characteristics**

| Subject | Age/Sex | Braak and Braak grade, duration | Cerebral amyloid angiopathy | Cause of death | Family history of dementia | Major comorbidities |
|---|---|---|---|---|---|---|
| AD1 | 64 / M | VI, 7 years | Mild | Pneumonia | No (PS1 -) | None |
| AD2 | 79 / M | VI, 10 years | None | Stroke | No | Diabetes, papillary thyroid carcinoma, atherosclerosis |
| AD3 | 81 / F | VI, 7 years | Mild | Pneumonia | 2 siblings | Atrial fibrillation |
| AD4 | 82 / F | IV, 5 years | None | Pneumonia | No | None |
| AD5 | 93 / M | III, 3 years | Severe | Pneumonia | No | None |
| AD6 | 62 / M | I, none | Severe | Intracerebral hemorrhage | No | None |
| AD7 | 86 / M | VI, unknown (longstanding) | Severe | Intracerebral hemorrhage | No | Hypertension |
| AD8 | 71 / M | VI, 7 years | Severe | Intracerebral hemorrhage | No | Waldenstrom's macroglobulinemia, diabetes |
| Control 1 | 27 / M | None | None | Congenital heart disease | No | None |
| Control 2 | 29 / F | None | None | Arrhythmia | No | None |
| Control 3 | 45 / M | None | None | Basilar artery occlusion | No | Atherosclerosis, hypertension, diabetes |
| Control 4 | 57 / F | None | None | Sepsis | No | Perforated colon |
| Control 5 | 62 / M | None | None | Sepsis | No | C.dificil colitis, liver transplant recipient |
| Control 6 | 69 / F | None | None | Heart failure | No | Dilated cardiomyopathy |
| Control 7 | 77 / M | None | None | Sepsis | No | Lymphoma, neutropenia, atrial fibrillation. |
| Control 8 | 80 / F | None | Mild | Ischemic heart disease | No | Atherosclerosis, neuroendocrine lung tumor |

**Supplemental Figure 1.** PLD3 mRNA levels do not correlate with cerebral atherosclerotic Disease

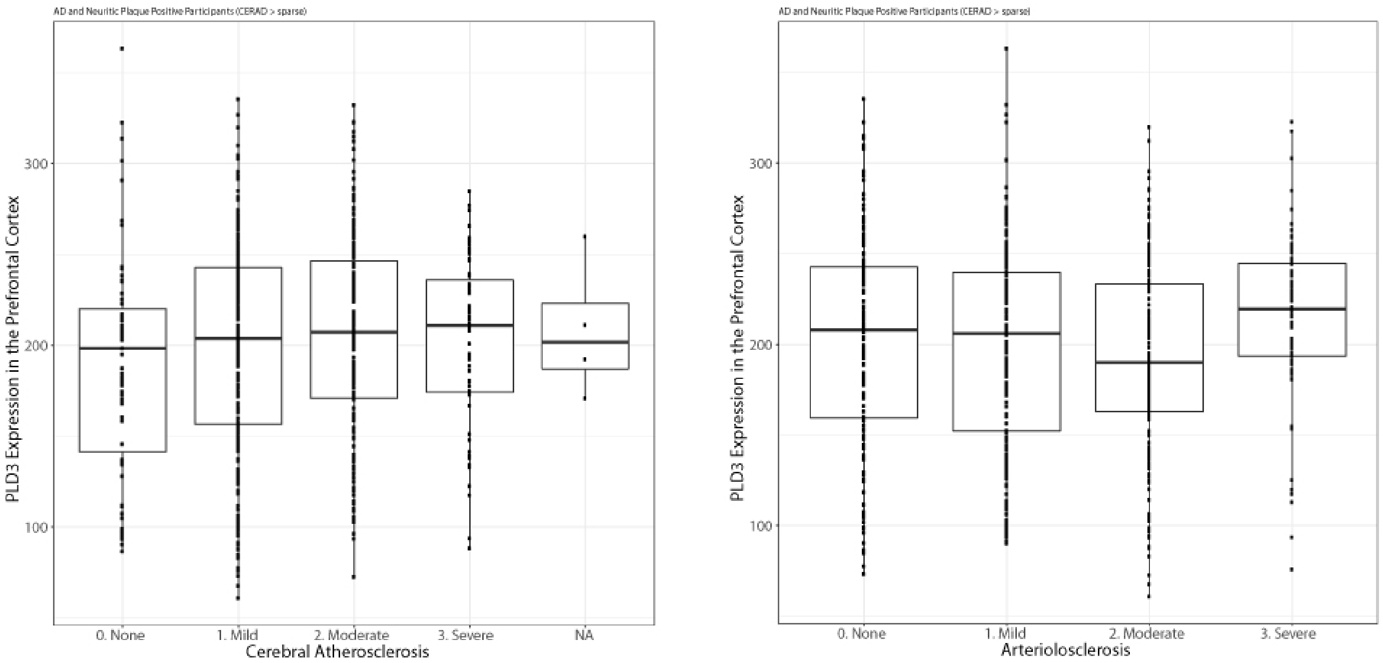

### Supplemental Figure 2. ROS/MAP Cognitive Sub-Measures

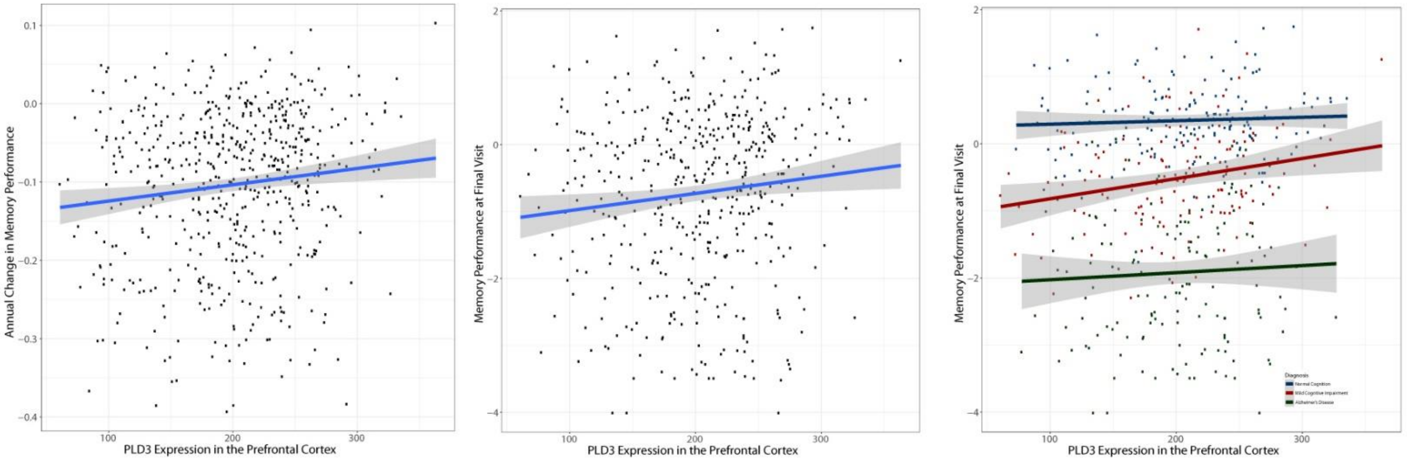

**Supplemental Figure 3: Protective effect of PLD3 expression on β-Amyloid burden is stronger in women**

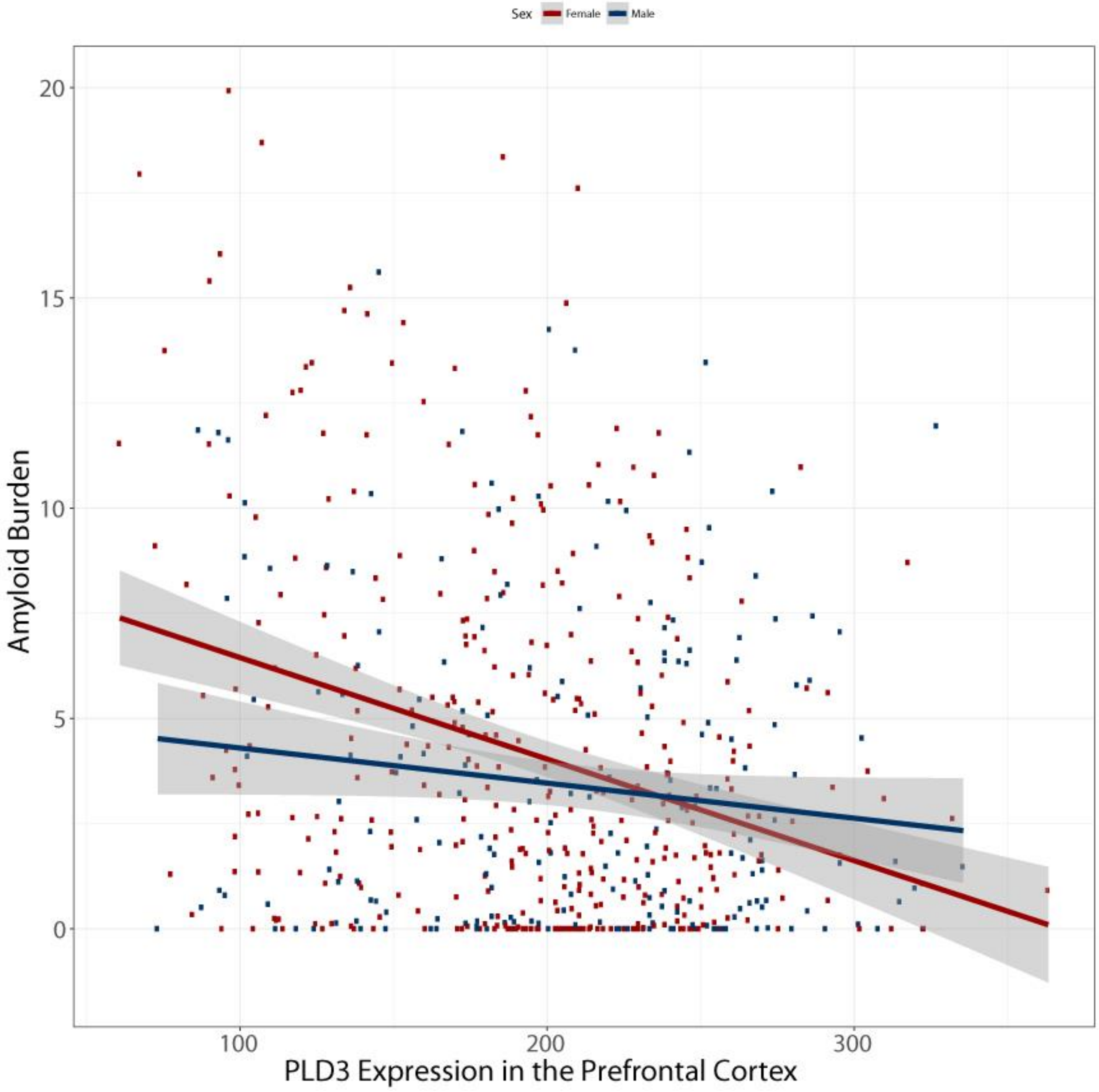

**Supplemental Figure 4.** Multiple dataset analysis reveals PLD3 signal in AD risk models

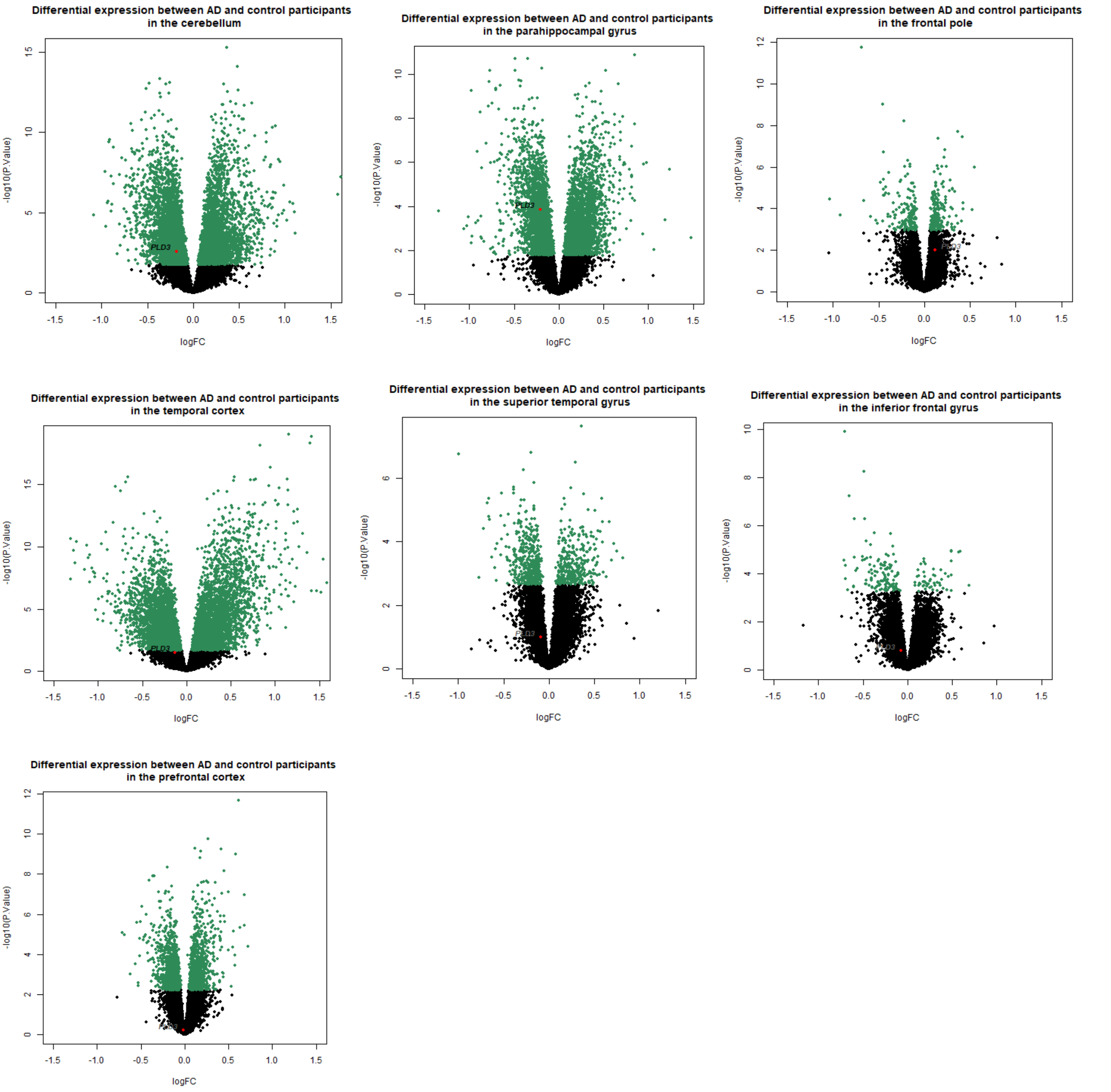


Volcano plots showing differential expression of all measured genes between AD participants and controls, highlighting *PLD3*. Green dots are genes which were significantly differentially expressed after correction for all genes tested in that tissue (17012 for Mayo tissues, 16348 for MSSM, and 15584 for ROSMAP).

**Supplemental Figure 5.** AD-BXD mouse performance on balance beam does not correlate with PLD3 expression

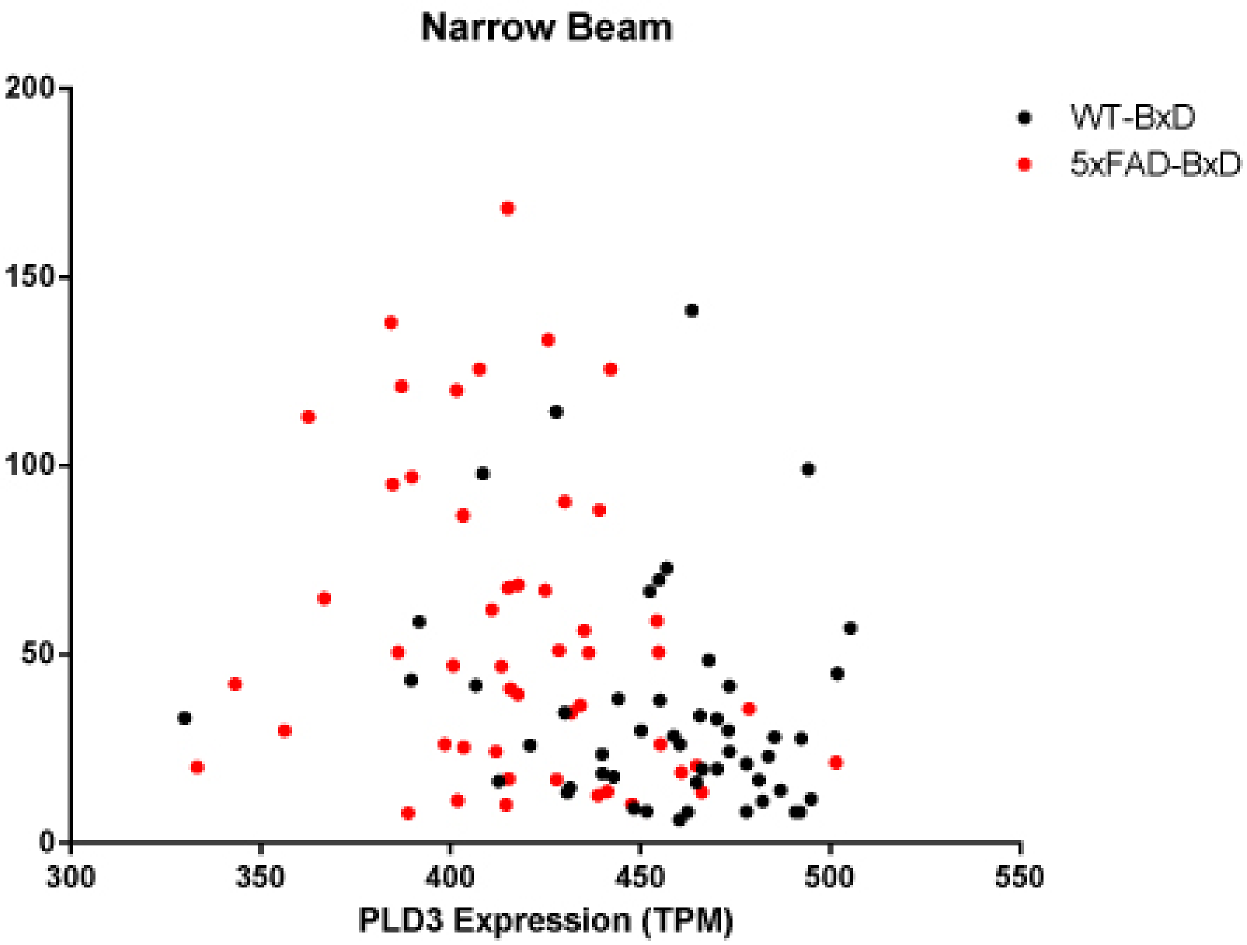


Narrow beam performance plotted against normalized PLD3 mRNA expression. Data collected as described previously[17]. PLD3 expression does not correlate with narrow beam performance, indicating that summated lack of PLD3 function does not induce motor deficits in this model. Linear regression reveals the plotted slope is not statistically non-zero ($p>0.1$), and this effect does not change upon the introduction of the 5xFAD transgene with respect to slope ($p=0.475$) or Y-intercept ($p=0.106$).

**Supplemental Figure 6: PLD3 colocalizes with numerous markers of lysosomes in human brain**

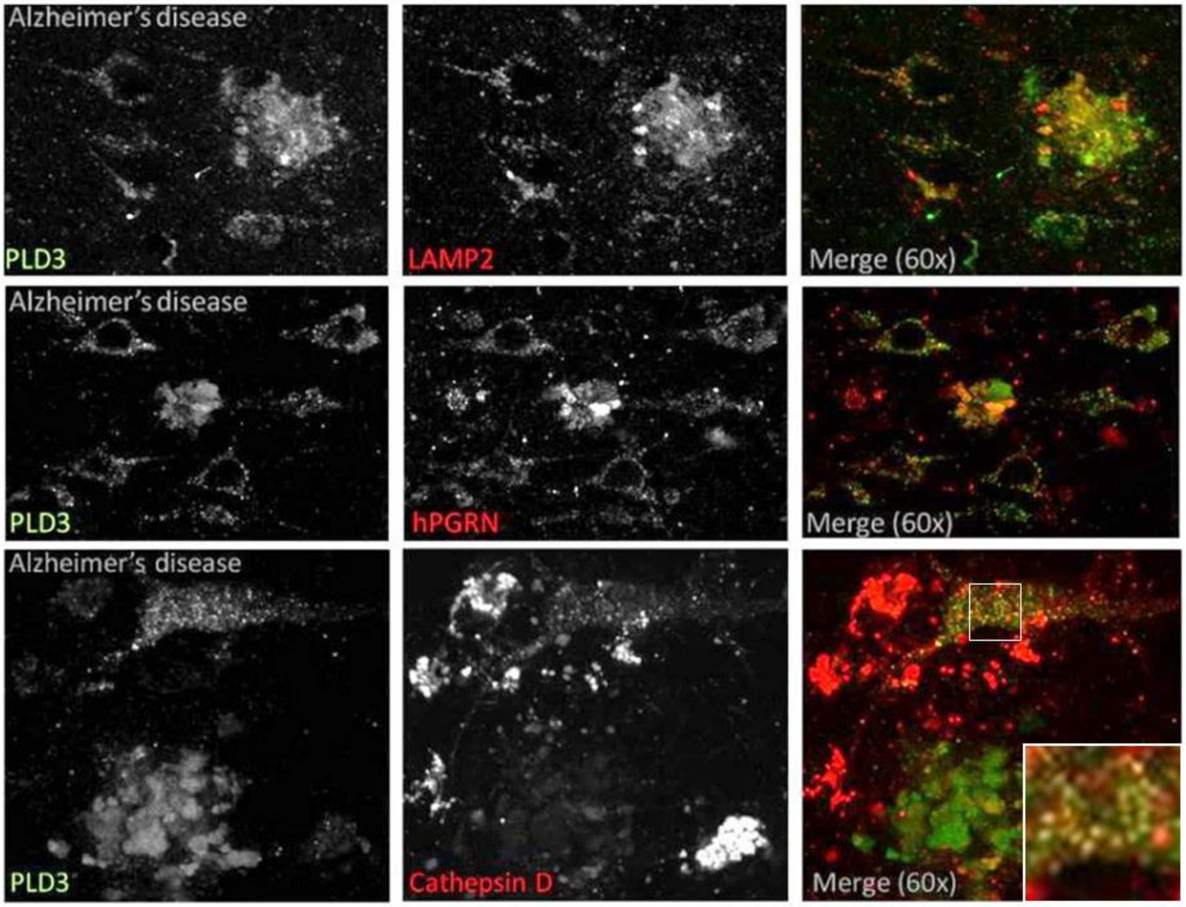


PLD3 is associated with lysosomes in human brain and massively enriched on abnormal lysosomes in dystrophic neurites. PLD3 closely colocalized with neuronal lysosomes and co-labeled with cathepsin D which is a luminal lysosomal protease, as well as LAMP2 and progranulin with are components of the lysosomal membrane. PLD3 accumulations around β-amyloid plaques co-label with LAMP2 and progranulin. Lysosomes in dystrophic neurites are deficient in cathepsins as has been previously reported in animal models of Alzheimer's disease.

### Supplemental Figure 7: Lysosome purification procedure validation

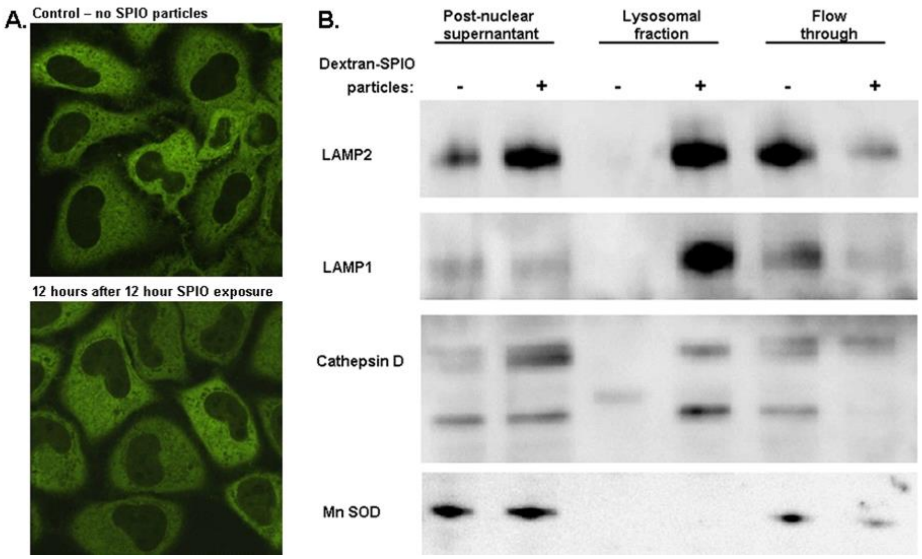


A. HeLa cells stably transfected with GFP-tagged transcription factor EB (TFEB), a transcription factor central to the initiation of lysosomal biogenesis, demonstrate cytosolic localization of TFEB in basal conditions. Upon induction of lysosomal biogenesis, TFEB rapidly translocates to the nucleus. To ensure that loading of dextran-coated small paramagnetic iron oxide nanoparticles (SPIO) did not induce lysosomal biogenesis, we demonstrated that 12-hour incubation with the nanoparticles did not induce TFEB translocation. B. Efficiency and purity of lysosome enrichment of lysosome isolate by magnetic chromatography are demonstrated by the enrichment of lysosome markers LAMP1, LAMP2 and Cathepsin D in the lysosomal fraction, without enrichment of mitochondrial marker manganese superoxide dismutase (Mn SOD).

**Supplemental Figure 8: Anti-PLD3 siRNA Depletes PLD3 levels**

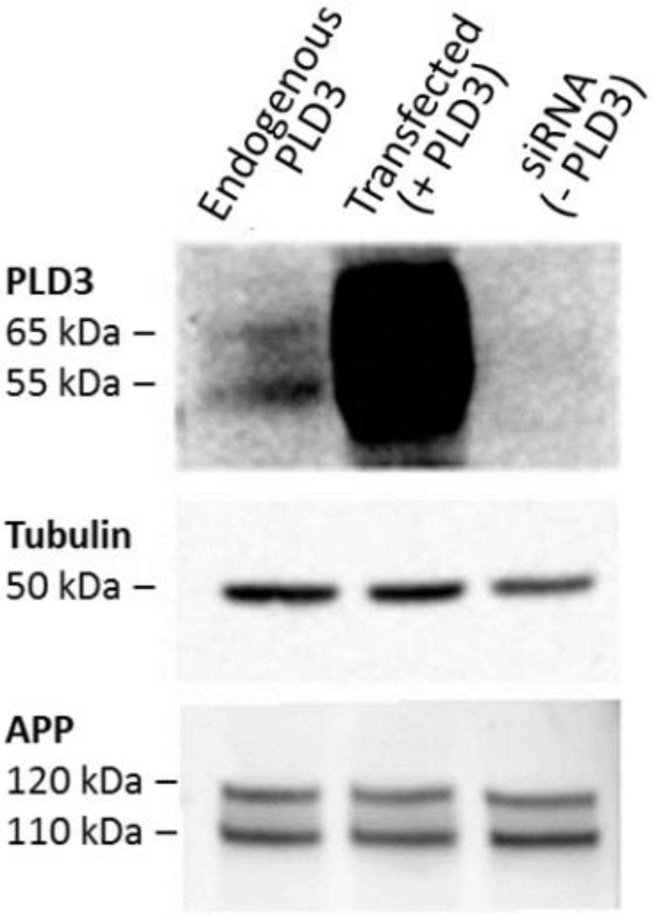


Endogenous PLD3 was detected in HeLa cells as a pair of bands on western blot at 55 and 65 kDa. These bands were eliminated by transfection of an siRNA against the human PLD3 transcript and amplified many fold by transfection of a plasmid containing an untagged PLD3 transcript. Manipulating the level of PLD3 did not alter the level of APP, confirming a previous report[7].

**Supplemental Figure 9: HeLa Derived Transfected Lysosomes Replicate PLD3 Specific Enzyme Function**

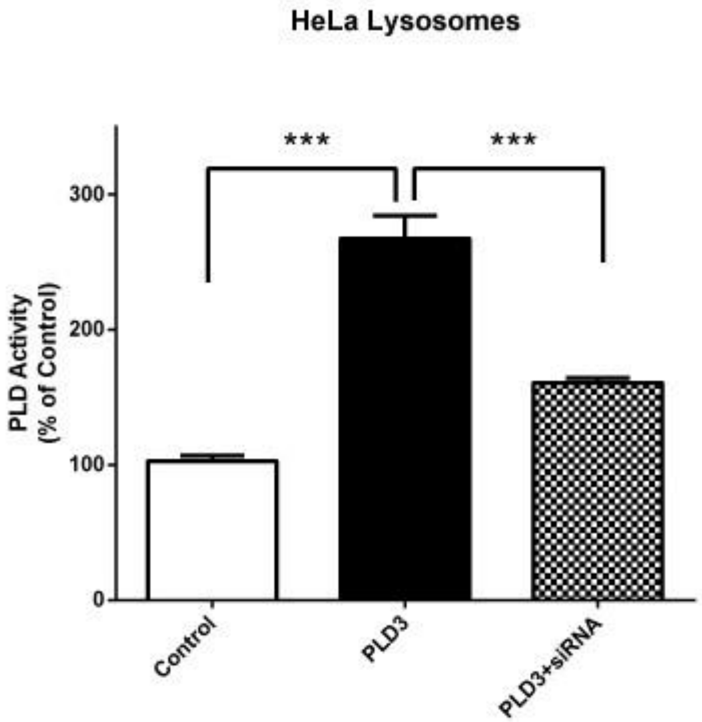


Replication study in HeLa cells reveals that PLD3 has enzymatic PLD activity in human cell line derived lysosomes. Lysosomes isolated from PLD3 transfected HeLa cells display significantly increased PLD activity compared to control lysosomes ($p<0.0001$). Cotransfection of PLD3 and siRNA against PLD3 significantly reduces lysosomal PLD activity ($p<0.0001$), indicating the specificity of the enzymatic activity of PLD3. PLD activity in PLD3+siRNA compared to control is not significant ($p=0.48$).

**Supplemental Figure 10: PLD3 transfection does not alter lysosomal level of PLD1 or PLD2**

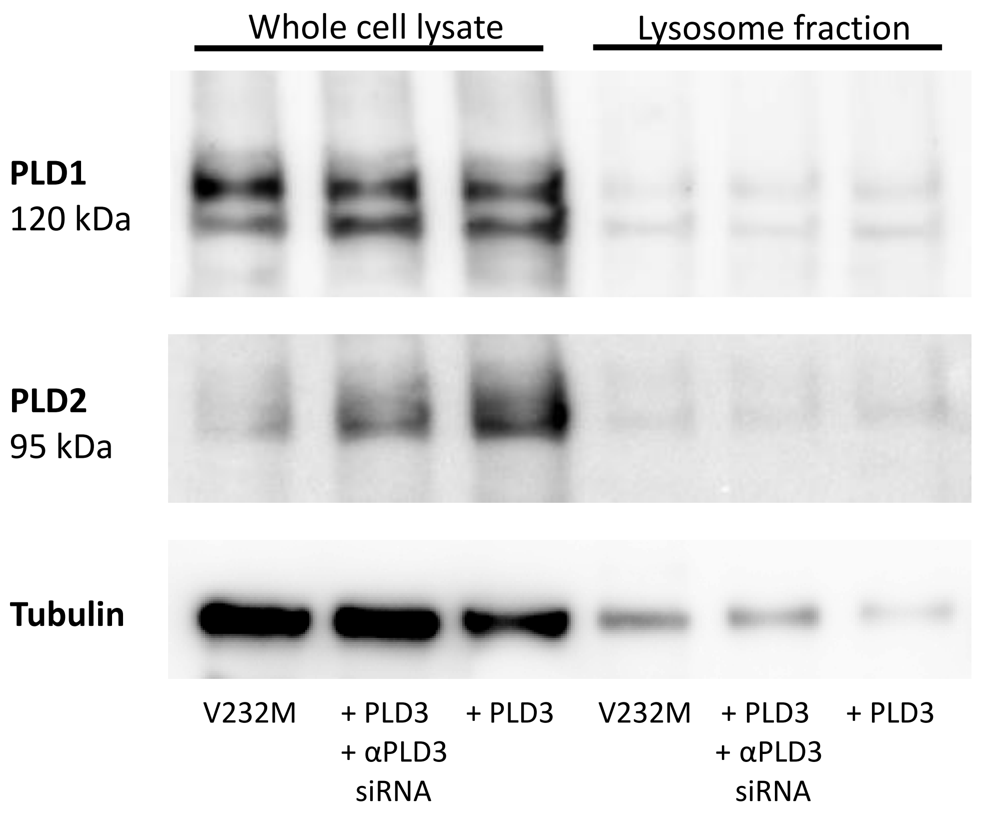

### Methods

#### Immunohistochemistry

*Human AD Brain Samples*

Formalin fixed brain tissue from eight human participants was obtained from autopsy, including eight cases of autopsy-verified Alzheimer's disease and eight neurological controls (**Supplementary Table 1)**. Tissue procurement was approved by the Institutional Review Board and all specimens were de-identified.

*Mouse Brain Samples*

WT and 5xFAD 3-month-old mice were euthanized following ketamine/xylazine anesthesia and transcardial perfusion with ice cold PBS. One hemisphere of each mouse was used for immunohistochemistry and was drop fixed in ice-cold 4% paraformaldehyde/PBS. Brains were left to fix for at least one week at 4°C before slicing.

*Brain Sectioning & Staining*

Fixed brain tissue was sectioned at 50 µM on a Leica vibratome. Sections were then subjected to antigen retrieval, consisting of firstly heating to 95°C in citric acid buffer (10mM citric acid, 0.05% Tween-20, pH 6) for 20 min. Slices were then transferred to PBS/glycine buffer (100 mM glycine/1x PBS, 0.1% Tween-20) for 10 min shaking at room temperature. Following that, slices were then washed in PBS/0.1% Triton X-100 for 10 min at room temperature. Sections were then placed in blocking buffer (4% Bovine serum albumin (BSA) in PBS/0.1% Triton X-100) under a 240-watt LED panel with emission at 390nm, 430nm, 460nm, 630nm, 660 nm and 850 nm (GrowLight, HTG Supply Inc) for a minimum of 18 hours at 4°C to reduce autofluorescence related to lipofuscin. Primary antibodies were applied in 4% BSA in PBS/0.1% Triton X-100 overnight at 4°C with gentle shaking. Slices were then washed 3 x 10 min room temperature in PBS/T before application of fluorescent secondary antibody in 4% BSA/PBS/T for a minimum of 2 hours room temperature incubation. Slices were finally washed 3 x 10 min room temperature in PBS/T before slide mounting and coverslipping.

*Microscopy*

Tissue was visualized on a laser scanning confocal microscope (Zeiss, LSM 710) with a 20x objective or 63x oil immersion objective. Images were acquired with a minimum resolution of at least 1024x1024 pixels. All images underwent routine processing with the Image J software (https://imagej.nih.gov/ij/download.html). Colocalization was assessed with Pearson's correlation coefficient and analyzed with MIPAV image analysis software (https://mipav.cit.nih.gov/download.php).

#### AD-BXD Studies

*AD-BXD Mouse Line*

Mice were generated as previously described[17,20]. Briefly, stable inbred BXD mouse lines (generating an array of genetic diversity between C57BL/6 and DBA2/J) were crossed with 5xFAD hemizygous mice, which contain 5 induced variants known to cause familial AD in human[32]. Due to the hemizygous nature of 5xFAD, 50% of the F1 progeny carry the 5xFAD insert (AD-BXD) and 50% are WT for littermate controls (WT-BXD), where all mice contain the respective BXD line gene diversity. Experiments were performed at 6 months and 14 months of age upon both male and female mice. Mice underwent contextual fear conditioning (n=636), of which a subset was euthanized and brain tissue subjected to RNA-Seq (n=133). Dots represent averaged behavioral performance within BXD background strain and PLD3 transcript count (n=100 observations).

*Contextual Fear Conditioning*

Following 3 days of habituation to transport and to the testing environment, mice were trained on a standard contextual fear conditioning paradigm as previously described[17,20]. Contextual fear training consisted of a 180s baseline period followed by four mild foot shocks (1s, 0.9mA), separated by 115±20s. A 40s interval following each foot shock was defined as the post-shock interval, and the percentage of time spent freezing during each of these intervals was measured using FreezeFrame software (Coulbourn Instruments, PA, USA). To calculate an acquisition curve for each strain as an index of learning, the slope across the average time spent freezing during post-shock intervals 1-4 was derived. Twenty-four hours later, hippocampus-dependent contextual fear memory (CFM) was tested by returning the mouse to the testing chamber for 10 min. The percentage of time spent freezing during the testing trial was measured using FreezeFrame software and used as an index of CFM.

*RNA-Seq*

RNA was quantified via RNA-Seq as previously described[17], and available on GEO (https://www.ncbi.nlm.nih.gov/geo/query/acc.cgi?acc=GSE101144). Briefly, snap frozen hippocampi from AD-BXD strains and non-carrier littermate controls at 6 and 14m were used for RNA sequencing. RNA was isolated using the RNeasy mini kit (Qiagen) and treated with DNase to remove contaminating DNA. Final library pools were sequenced by 75bp paired-end sequencing on a HiSeq2500 (Illumina Inc). For final by-strain analysis, samples belonging to the same strain/sex/age/genotype group were averaged.

**ROS-MAP Cohort and Analysis**

Data for human analyses were acquired from two cohort studies acquired and made available by the Rush University AD center (RADC), all data of which are freely available through the RADC web portal (www.radc.rush.edu). The Religious Orders Study (ROS) began in 1994 and the Rush Memory and Aging Project (MAP) began in 1997. Both studies enrolled older adults without dementia and all participants signed an Anatomical Gift act for brain donation[13–15]. Written informed consent was obtained from all participants and all protocols were approved by the Institutional Review Board (IRB). Secondary analyses of data were approved by the Vanderbilt University Medical Center Institutional Review Board.

*Clinical Diagnosis*

As described in detail previously[33], all clinical and neuropsychological data were reviewed by a board-certified neurologist at the time of death and a summary diagnostic opinion was made blinded to all postmortem data. Consensus diagnosis was reached in case conference for selected cases.

*RNA Sequencing*

RNA expression levels were obtained, quantified, processed, and normalized previously, and have been described in detail elsewhere[16,34]. Briefly, RNA expression levels were obtained from frozen, manually dissected sections of dorsolateral prefrontal cortex. RNA was isolated using the RNeasy lipid tissue kit (Qiagen, Valencia, CA) and was reverse transcribed and biotin-UTP labeled using the llumina® TotalPrep™ RNA Amplification Kit from Ambion™ (Illumina, San Diego, CA). Expression signals were generated using BeadStudio (Illumina, San Diego, CA). Standard control and normalization methods were employed to account for technical variability due to differences in hybridization dates. Reads were aligned using the Bowtie 1 package and then counted using RSEM, as previously described[34].

*Neuropathology*

Measures of AD neuropathology were previously quantified and described in detail elsewhere[13,14,34]. Amyloid and tau burden were quantified in eight brain regions using immunohistochemistry including the superior frontal cortex, anterior cingulate, hippocampus, angular gyrus, entorhinal cortex, calcarine cortex, middle frontal cortex, and inferior temporal cortex. Neuritic plaques and neurofibrillary tangles were quantified from silver stained slides, including five brain regions (middle frontal cortex, middle temporal cortex, inferior parietal cortex, hippocampus, and entorhinal cortex). Each AD neuropathology outcome was square-root transformed to better approximate a normal distribution.

In addition, non-AD pathologies were previously quantified and described in detail elsewhere[35–41] including TDP-43[42], cerebral amyloid angiopathy (CAA)[37], microscopic infarcts[36], gross infarcts[40], arteriolosclerosis[38], and atherosclerosis[35]. The quantification and modeling of each of these outcomes for the present analytical models has been described previously[34].

*Cognitive Outcomes*

Cognitive performance was evaluated with 19 neuropsychological tests as previously described[43]. Tests probed multiple cognitive domains, including semantic memory, working memory, episodic memory, perceptual speed, and perceptual organization. Z-scores from all tests were averaged into a single global cognitive composite as previously described[43].

*Statistical Analyses*

All statistical analyses were performed in R (version 3.5.0; https://www.r-project.org/). Differential expression of *PLD3* across diagnostic groups was assessed using linear regression with *PLD3* entered as a continuous outcome and diagnosis as a categorical predictor with the normal cognition group set as the referent. Linear regression models assessed the association between *PLD3* and neuropathological burden at autopsy with square-root transformed AD neuropathology traits set as continuous outcomes in separate models. In all models, covariates included age at death, sex, and post mortem interval.

For cognitive analyses, linear regression models assessed the association between *PLD3* expression and cognitive performance at last visit prior to death in the same manner outlined above. Longitudinal change in cognition was evaluated using mixed-effects regression models with the intercept and interval (years from death) entered as both fixed and random effects and included the same covariates as the cross-sectional models. PLD3 associations with longitudinal change were assessed with a *PLD3* x interval interaction term in the model.

#### Accelerating Medicine Partnerships Alzheimer's Disease (AMP-AD) Replication Analyses

As described previously[34], harmonized data from the Mount Sinai Brain Bank (syn3159438) and the Mayo Clinic brain bank (syn5550404) were evaluated for replication leveraging resources from AMP-AD. Mount Sinai data included quantification of gene expression in the frontal pole, superior temporal gyrus, inferior temporal gyrus, and parahippocampal gyrus[44]. The Mayo data included expression from the cerebellum and temporal cortex[45,46]. Differential expression analyses for all measured genes were completed previously and used for replication in the present manuscript.

#### Cell Culture and PLD3 Expression

HeLa and NSC34 cells were cultured at 37ºC and 5% $CO_2$ in DMEM cell media containing 10% fetal bovine serum and penicillin and streptomycin. Cells were plated onto 150mm culture flasks and transfected with 20µg of pCMV3-hPLD3 (HG16349-NH; Sino Biological, Wayne, PA, USA) utilizing Fugene 6 (Promega, USA) at ~75% confluency. siRNA was transfected using Oligofectamine (Invitrogen). Target specificity of siRNA and efficiency were confirmed by western blot (**Supplemental Figure 1**). Experiments were performed 48h after transfection.

#### Lysosomal Isolation and Enzymatic Assay

Lysosomes were isolated similarly to procedures previously described[47]. Briefly, cells were grown to confluence on a 150mm dish and treated for 2 hours with a colloid suspension of dextran-coated iron oxide nanoparticles at a concentration of 10mg/mL in DMEM containing 10% fetal bovine serum and penicillin and streptomycin. The cells were then washed three times and allowed to rest for at least 2 hours prior to collection. Cells were scraped free in 1mL cold homogenization buffer consisting of PBS containing a protease inhibitor cocktail. The material was homogenized with 8 strokes of a tight fitting Dounce homogenizer and was applied to a column containing 50mg fine steel filaments which was placed on a strong magnet. The cellular material passing through the column was collected and applied to the column two more times. The column was then washed with 10mL cold homogenization buffer. The column was then placed in a microcentrifuge tube and centrifuged at 10,000g for 2 minutes yielding the lysosomal fraction. Approximately 1.5-3% of total protein was recovered in this fraction. The isolation procedure was conducted in less than 1 hour and all steps were performed on ice. Protein concentration was assessed via the Bradford method (reagent). Equal protein of intact lysosomes was loaded in replicate onto 96-well plates and then incubated with Amplex Red PLD assay kit, per manufacturer instructions. Samples were incubated at 37°C for 1 hour, and read on a fluorescent plate reader (POLARstar Omega; BMG Labtech, USA) at (Excitation: 571nm/Emission: 585nm)

**Western blot**

Cells were scraped from 150mm dishes in 1mL cold PBS containing 1% Triton X-100 and a protease inhibitor cocktail. Cells were rotated end-over-end at 4°C for 60 minutes, then centrifuged at 10,000g for 10 minutes to remove crude cellular debris. Protein concentration of the resulting lysate was determined with a Bradford assay. The lysate was combined with an equal volume of 2x Laemmelli's buffer containing beta-mercaptoethanol and heated to 95°C for 10 minutes to denature. Equal protein mass was loaded into each lane of a pre-cast 4-15% gradient acrylamide gel and run at 200V. Transfer was performed in a semi-dry apparatus (Trans-Turbo, BioRad). PVDF membrane were then blocked in 4% bovine serum albumin (BSA) in phosphate buffered saline (PBS) containing 0.1% Tween-20 (PBST) for 60 minutes. They were then incubated with primary antibodies diluted in 4% BSA in PBST overnight. Membranes were then washed 3 x 10 min in PBST prior to incubation with secondary antibody (diluted in 4% BSA in PBST) for 2 hours. Membranes were washed three times again, treated with Clarity chemiluminescence reagent and developed on a chemiluminescence imaging system (ImageQuant LAS 4000)

**Graphical and Statistical Methods.** All data were graphed and statistically analyzed using Prism 6.0 (GraphPad, Inc., La Jolla, CA) or R, as noted. In all statistical tests, P value significance thresholds were set *a priori* to $\alpha=0.05$. Specific statistical tests to assess main effects and post-hoc tests as well as N for all groups are provided in the Figure Legends.
